# Measuring diversity from space: a global view of the free and open source rasterdiv R package under a coding perspective

**DOI:** 10.1101/2020.11.14.369371

**Authors:** Elisa Thouverai, Matteo Marcantonio, Giovanni Bacaro, Daniele Da Re, Martina Iannacito, Carlo Ricotta, Clara Tattoni, Saverio Vicario, Duccio Rocchini

## Abstract

The variation of species diversity over space and time has been widely recognised as a key challenge in ecology. However, measuring species diversity over large areas might be difficult for logistic reasons related to both time and cost savings for sampling, as well as accessibility of remote ecosystems. In this paper, we present a new `R` package - `rasterdiv` - to calculate diversity indices based on remotely sensed data, by discussing the theory beyond the developed algorithms. Obviously, measures of diversity from space should not be viewed as a replacement of in-situ data on biological diversity, but they are rather complementary to existing data and approaches. In practice, they integrate available information of Earth surface properties, including aspects of functional (structural, biophysical and biochemical), taxonomic, phylogenetic and genetic diversity. Making use of the `rasterdiv` package can result useful in making multiple calculations based on reproducible open source algorithms, robustly rooted in Information Theory.

## 1 Introduction

Back in 1872, Ludwig Eduard Boltzmann (Boltzmann, 1872) introduced the first measure of entropy, later called marginal entropy and restructured by Claude Elwood Shannon under a mathematical theory umbrella (Shannon, 1948). As such, it became one of the cornerstones of ecological theory and was adopted widely in ecological practice for measuring biodiversity and its change. Concerning biological entropy, the variation of species diversity over space and time has been widely recognized as a key challenge in ecology and was associated with analytic geometric models focusing either on the spatial component of species dispersal (Palmer, 2007; Gorelick, 2008) or on environmental drivers (Kreft and Jetz, 2007).

To address this issue, many spatio-statistical models have been proposed to model biological entropy using data from ecological surveys (Bachl et al., 2019). However, the statistical clarity (sensu Dushoff et al. (2019)) of such models strictly depends on a high in-situ data uncertainty, which propagates through all inferential steps (Meyer et al., 2016; Rocchini et al., 2019). Furthermore, measuring species diversity over wide areas might be difficult for logistic reasons related both to time and sampling costs (Chiarucci et al., 2011) and to theoretical and practical constraints, which are mainly related to two sources of uncertainty. The first is the uncertainty associated with the detectability and the determination of individual plants or animals species. The second is the one linked to different sampling strategies (McGlinn and Palmer, 2009) or efforts (Rocchini et al., 2019) per area, or, in the worst case, to the impossibility of getting information about the real grain (sensu Scheiner et al. (2000)) sampled (Hobohm, 2003). In the absence of such information, it becomes excessively challenging to properly address the modifiable areal unit problem (MAUP), which in this case is the sensitivity of biodiversity to scale (Jelinski and Wu, 1996). This is true, even though evidence exists for a chance to rely, in some instances, on expert knowledge to build straightforward and robust diversity maps worldwide (Hobohm et al., 2019).

Accordingly, algorithms based on remote sensing and spatial ecology might help estimating the variation of biodiversity over space and time (Skidmore et al., 2011; Schimel and Scheiner, 2019) and represent a powerful first exploratory tool to detect the spatial variability accross the landscape. The relationship between ecological processes (and functions) and the remotely sensed diversity can rely on the definition of niche proposed by Kroes (1977), and according to which a niche is the biotic structural and functional part of the ecosystem. Strictly speaking, such definition can be profitably used to measure spatial heterogeneity in ecosystems in order to convey information on their potential functions (Scheiner et al., 2017).

From this point of view, the development of Free and Open Source algorithms to measure diversity from space would be beneficial to allow high robustness and reproducibility of the proposed approaches (Rocchini and Neteler, 2012). Furthermore, their intrinsic transparency, community-vetoing options, sharing and rapid availability are also valuable additions and reasons to move to open source options. Among the different open source packages, the `R` software environment is certainly one of the most widespread world-wide and different packages have been devoted to remote sensing for: i) raster data management (`raster` package, Hijmans and van Etten (2020)), ii) remote sensing data analysis (`RStoolbox` package, Leutner et al. (2019)), iii) spectral species diversity (`biodivMapR` package, Féret and Boissieu (2020)), iv) Sparse Generalized Dissimilarity Modelling based on remote sensing data (`sgdm` package, Leitao et al. (2012)), v) entropy-based local spatial association (`ELSA` package, Naimi et al. (2019)), vi) landscape metrics calculation (`landscapemetrics` package, Hesselbarth et al. (2019)), to name just a few. Reader can also refer to https://cran.r-project.org/web/views/Spatial.html for the CRAN Task View on analysis of spatial data.

However, currently no package provides a flow of functions grounded on Information Theory related to abundance based measures, by further introducing distances and going back to Information Theory again by generalised entropy. In this paper we introduce a new R package which provides such a functions’ throughput workflow. The aim of this manuscript is to encompass the theory beyond the algorithms developed in the `rasterdiv` package (currently available at: https://github.com/mattmar/rasterdiv), relying on the definition given by Gorelick (2011b):

“*Theory is neither mathematical nor abstract. Theory is the creative, inductive, and synthetic discipline of forming hypothesis* […]”

## 2 Information Theory

One of the mostly used metrics for measuring remotely sensed diversity is related to the entropy measurement firstly introduced by Shannon (Shannon, 1948).

Given a sample area with *N* pixel values and *p*_*i*_ relative abundances for every *i* ∈ {1, …, *N*}, in decreasing order, the Shannon index is calculated as:

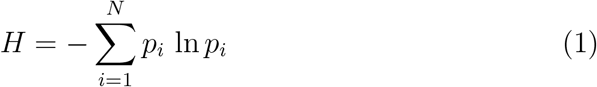

Taking into account only the most abundant pixel value, the Berger-Parker (Berger and Parker, 1970) index is given by:

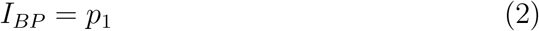

In remote sensing applications, the derivation of synthetic indices of any sort (i.e., diversity) is often performed by sequentially considering only small chunks of the whole image. These chunks are commonly defined as ’kernel’, ’windows’ or ’moving windows’. From now on, we will use this terminology to indicate the local space of analysis.

Both indices can be calculated in `rasterdiv` on a numerical matrix by using a moving window and applying the command `Shannon` and `BergerParker`.

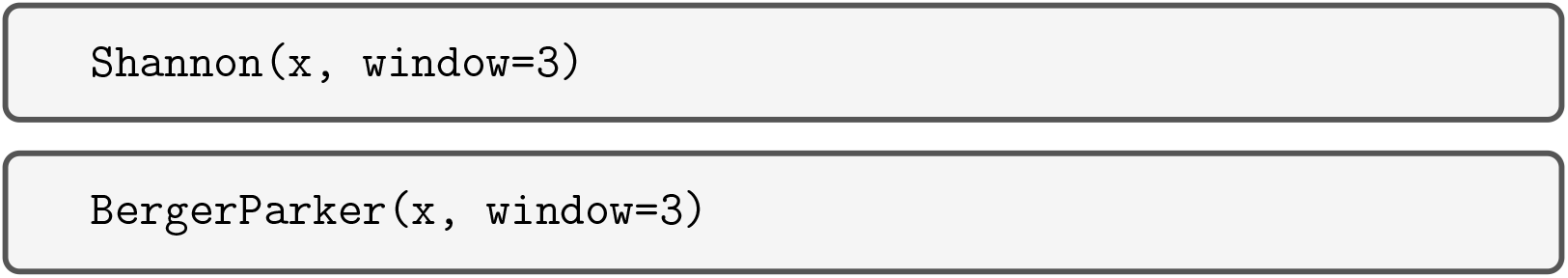

where `x` and `window` represent the input numerical matrix (raster file) and the side of the moving window on which the calculation is performed. Additional arguments common to all functions in the package are `np` which sets the number of parallel processes among which to split the calculation of the index, `cluster.type` which defines the type of protocol to spawn parallal processes (default is `‘‘MPI’’`) and `debugging` that, if set as `TRUE`, will run the function in verbose mode.

Both indices obey to the relative abundance of values. The most simple Berger-Parker index equals the relative proportion of the most abundant class in a moving window (Figure 1). Hence, low values of Berger-Parker are expected for continuous satellite data, given the high variability of reflectance values. In contrast, in the Shannon index, the abundance of every single numerical category (pixel value) is taken into account. This might lead to taking into account the turnover among values, since the higher the turnover the lower the dominance of a single class (Figure 2). However, Shannon’s *H* is unable to discern situations where there is a high richness (number of numerical categories) and a low evenness from those where there is a low richness but a high evenness.

**Figure 1:**
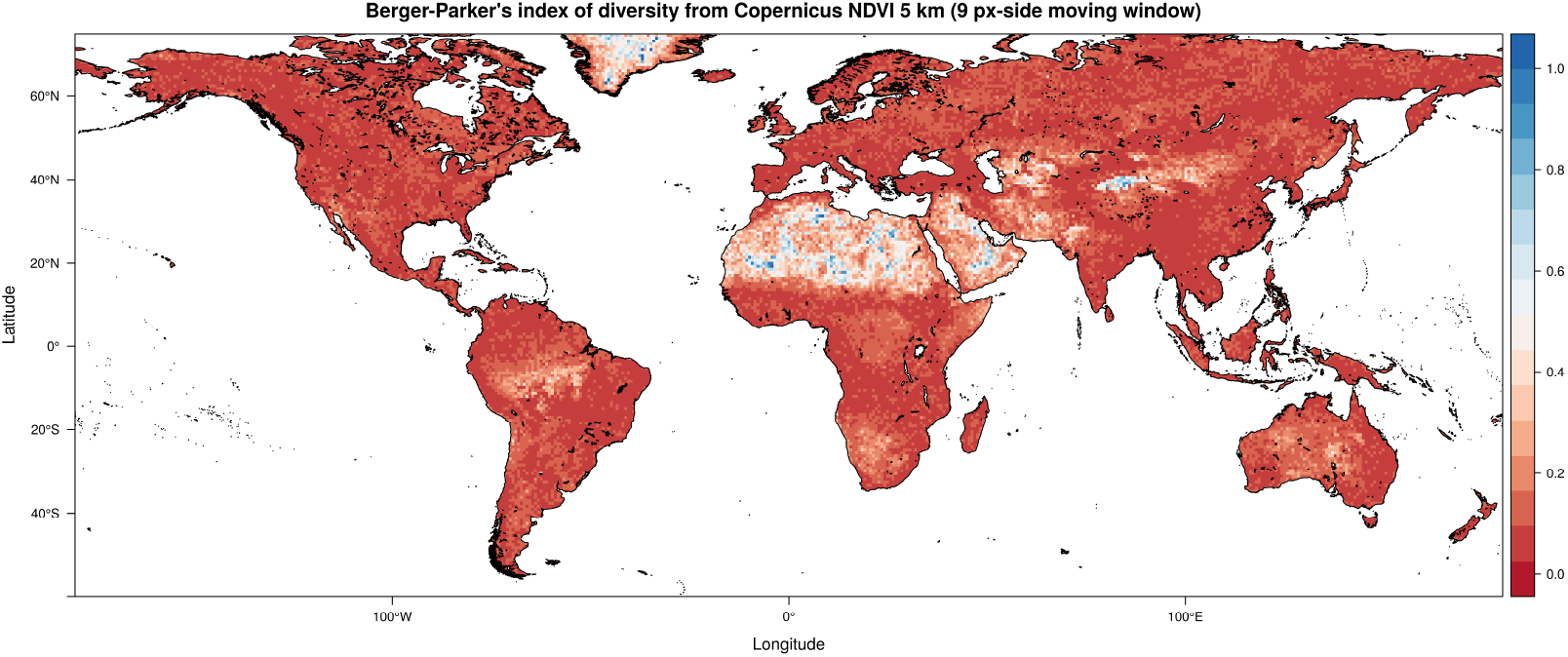
Berger-Parker index measuring the most abundant spectral value (Equation 2). All the indices in this paper are calculated starting from a Copernicus Proba-V NDVI (Normalised Difference Vegetation Index) long term average image (June 21st 1999-2017) at 5km grain, also provided into the `rasterdiv` package as a free default set which can be loaded by the function `data()`. A generally low value of the index (based on the most abundant spectral value) is found, since spectral input values are generally different from each other in a moving window. This figure has been generated by the command `bpa <-BergerParker(ndvi17 r,window=9,np=8,cluster.type=“SOCK”)`.

**Figure 2:**
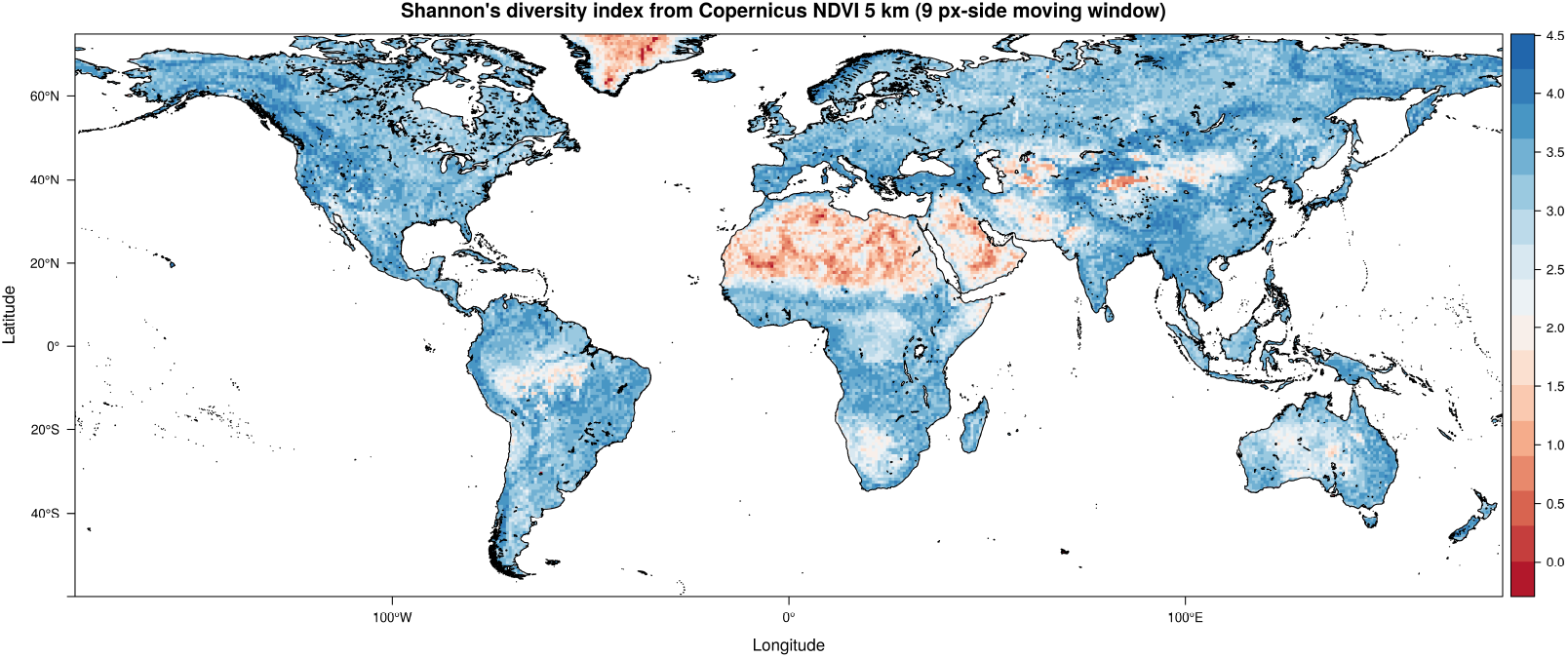
Shannon index calculated on a Copernicus Proba-V NDVI image at 5km. Shannon’s H is generally high since it only considers relative abundance of spectral values, which are generally different from each other. This figure has been generated by the command `sha <-Shannon(ndvi17 r,window=9,np=8,cluster.type=“SOCK”)`.

To better account for evenness, the Pielou index (Pielou, 1966) can be calculated by simply standardising the Shannon index on the maximum possible Shannon index attainable given the same richness value. The latter is attained when the maximum potential evenness of pixel values/numerical categories is reached, i.e. when they are equally distributed over space.

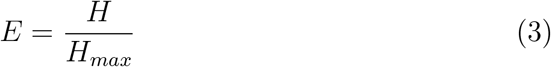

*H*_*max*_ corresponds to the natural logarithm of the number of pixel values. Using rasterdiv, the Pielou index can simply be calculated as:

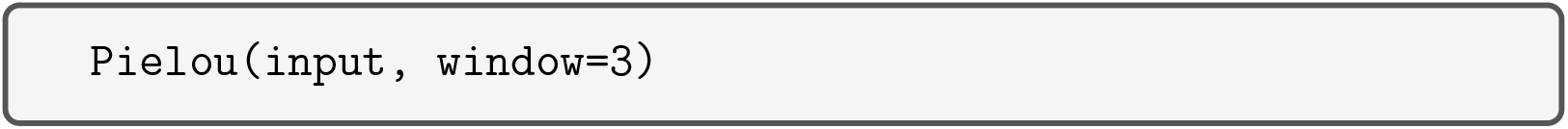

Figure 3 reports an example with a moving window of 9×9 pixels.

**Figure 3:**
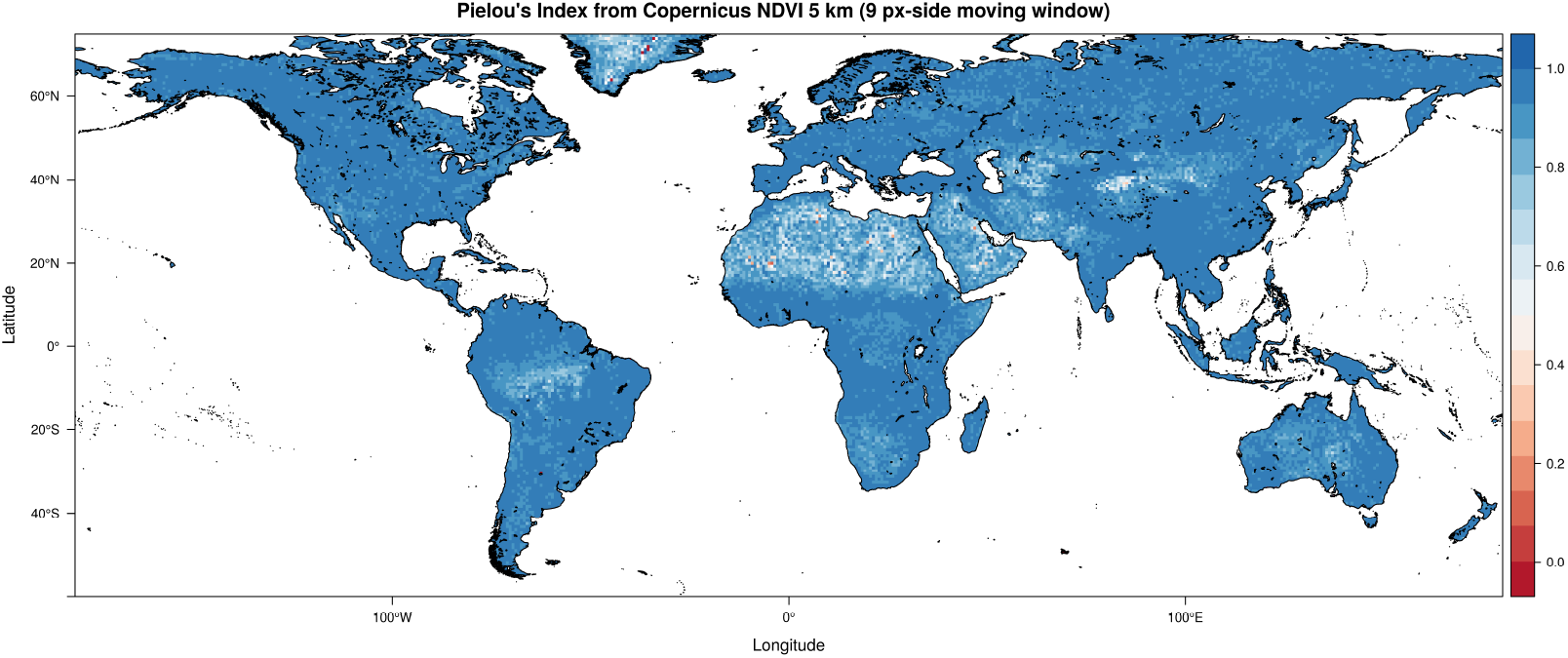
Pielou index calculated on a Copernicus Proba-V NDVI image at 5km. A flattening effect with respect to Shannon’s H is found, due to the standardisation on the maximum possible Shannon entropy (see Equation 3). This figure has been generated by the command `pie <-Pielou(ndvi17 r,window=9,np=8,cluster.type=“SOCK”)`.

## 3 Solving the non-dimensionality of Shannon’s *H*′: the Rao’s Q diversity index

Both Shannon’s *H* and Pielou’s *E* are dimensionless. In other words, they consider differences in the relative abundance among pixel values, but not their relative spectral distance, i.e. the distance among spectral values. For instance, let *A* = (1, 2, 3, 4, 5, 6, 7, 8, 9) and *B* = (1, 10^2^, 10^3^, 10^4^, 10^5^, 10^6^, 10^7^, 10^8^, 10^9^) be two theoretical arrays of values. In both cases, values are different from each other; hence, despite their relative numerical distance the Shannon index will always be maximum, i.e. *H* = log(9) = 2.197225 reducing *E* = *H/H*_*max*_ = 1.

In remotely sensed imagery this is a crucial point since it might happen that contiguous zones could have similar (but not equal) reflectance values. For instance, the diversity of a homogeneous surface like water could be overestimated if spectral distances are not considered.

To overcome this issue, the Rao’s Quadratic diversity (hereafter Rao’s Q, Rao (1982)) could be applied by not only taking into account relative abundance but also the spectral distance among different pixel values.

Given the values of different pixels *i* and *j*, the Rao’s Q consider their pairwaise distance *d*_*ij*_ as:

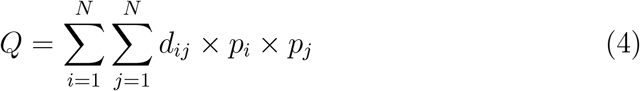

Hence, an array with different but spectrally close values will lead to a high Shannon’s H but a low Rao’s Q. On the contrary, an array with different and distant values in the spectral space will lead to both a high Shannon’s H and a high Raos’ Q.

Moving towards a 2D spatial extent, let M be a 2D matrix 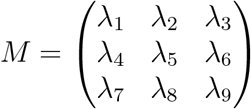formed by pixels with a certain reflectance value *λ* in a single band for instance. For simplicity, let us consider an 8-bit band, i.e. containing 256 possible values. As a consequence, deriving Rao’s Q involves calculating a distance matrix *M*_*d*_ for all the pixel values:

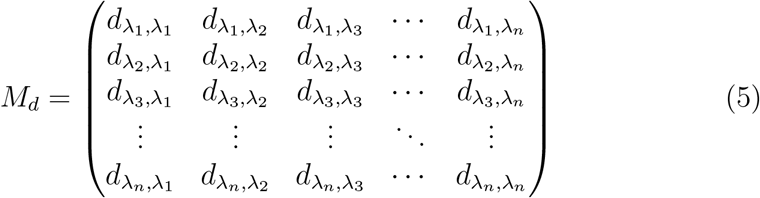

Thus, according to Equation 4, Rao’s Q is related to the sum of all the pixel values pairwise distances, each of which is multiplied by the relative abundance of each pair of pixels in the analysed image *d* × (1/*N*^2^). In other words, Rao’s Q is the expected difference in reflectance values between two pixels drawn randomly with replacement from the evaluated set of pixels. The distance matrix can be built in several dimensions (layers), thus allowing to consider more than one band at a time. As a consequence, Rao’s Q can be calculated in a multidimensional (multi-layers) system.

In `rasterdiv` package Rao’s Q is calculated as:

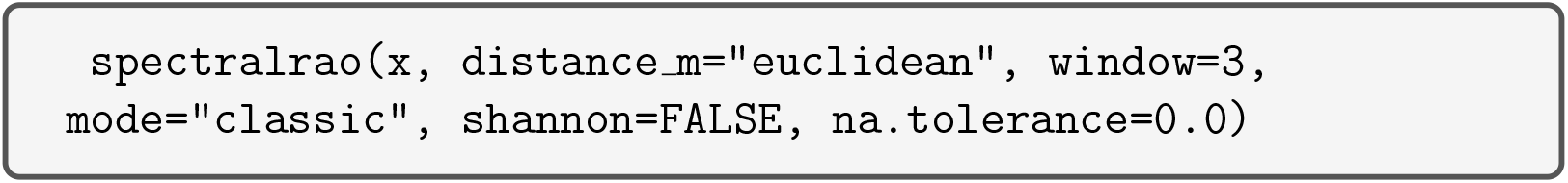

The `distance m` argument includes different types of distances, such as the Euclidean, Manhattan and Canberra distances, to make the calculation. Obviously, it is suggested to make use of a different distance when a multi-spectral set is used; otherwise all the considered distances in just one band will reduce to the Euclidean one (Figure 4).

**Figure 4:**
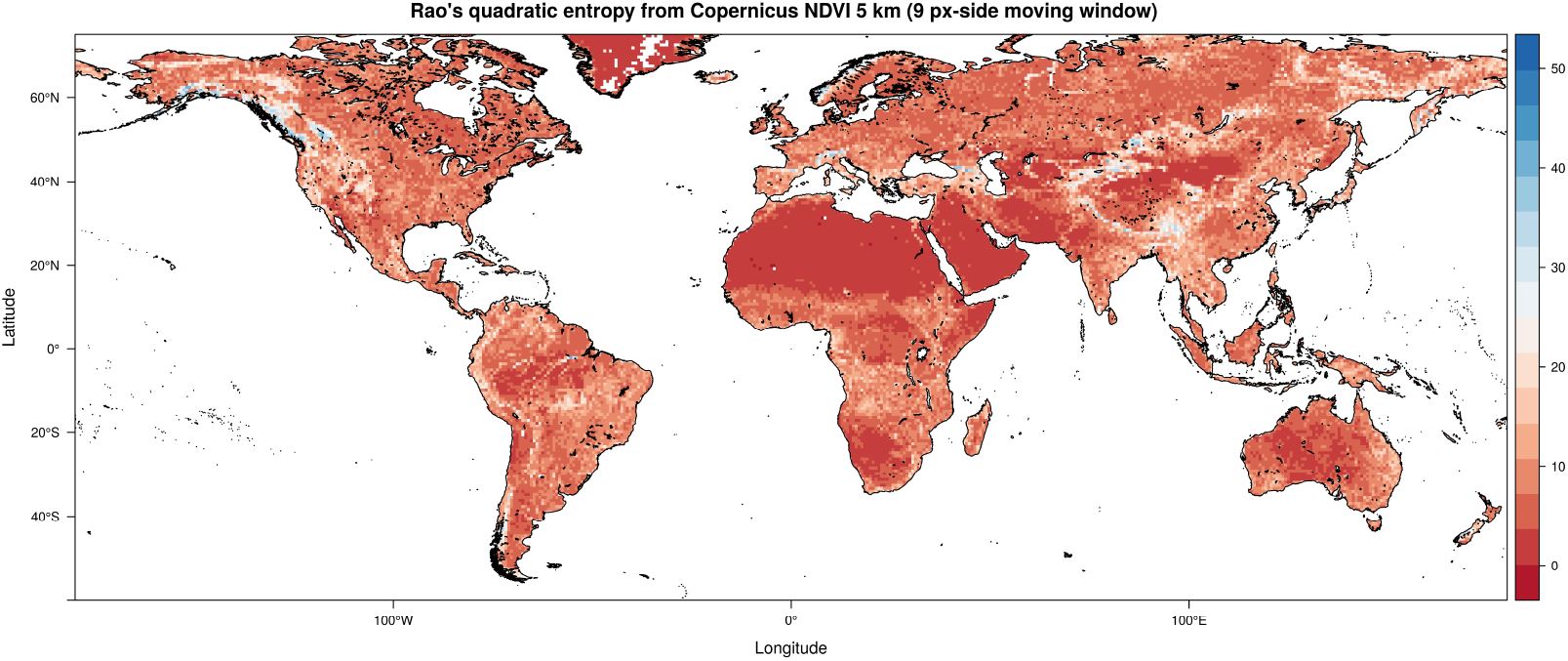
Rao’s Q index calculated on a Copernicus Proba-V NDVI image at 5km. Differenly from the original Shannon’s formula, Rao’s Q also considers the distance among different values by better discriminating the queues of the diversity distribution from very low diversity (e.g. deserts and ice fields) to very high diversity (e.g. upper highly-complex mountain ranges). This figure has been generated by the command `rao <-Rao(ndvi17 r,dist m=“euclidean”,window=9,shannon=FALSE,np=8, cluster.type=“SOCK”,na.tolerance=0.5)`.

In fact it is automatically demonstrated that in one dimension 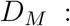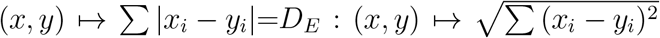 where *D*_*M*_ and *D*_*E*_ are the Manhattan and the Euclidean distances, respectively.

The Canberra distance is derived from the Manhattan distance by standardizing separately the absolute differences of each band with the sum of both values, and thus will also equal *D*_*E*_ in one dimension, such that: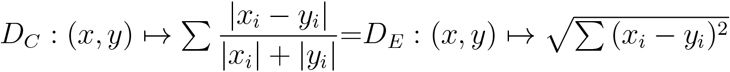.

## 4 Solving the intrinsic continuity of spectral data: Cumulative Residual Entropy

As previously stated, spectral data are continuous variables that are approximate to discrete (the so called “digital number”) for practical reasons. As such, the fact that two different pixels should be counted or not in a category depends from the whim of the normalisation of the signal when Digital Numbers (DNs) are generated. Shannon index is built strictly for a non ordered finite set of categories. For continuous variables, a derivative version of Shannon index was proposed, but soon it was clear that it had very different properties than categorical formulation (Jumarie, 1990; Michalowicz et al., 2013). Rao et al. (2004) proposed a Cumulative Residual Entropy (CRE) to build a consistent Shannon-like index for continuous variables. It is based on residual cumulative probability (*P* (*X* >= *x*_*i*_)), that can be estimated in a robust manner from empirical mono-dimensional distributions by counting for each value the number of observations with equal or larger values and then dividing by the total. CRE is defined as follow:

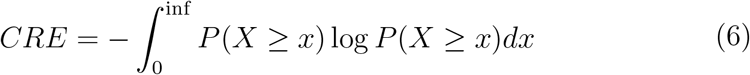

and to estimate it from an empirical distribution, the following approach is advised:

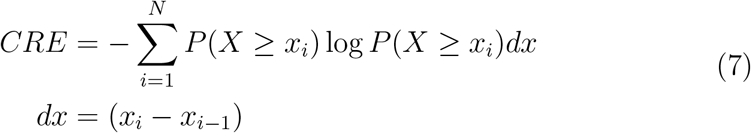

where *X* is the sorted vector of *N* observations. In practice, the approach is similar to the Rao’s Q, given that a coefficient *d*, representing the disparity of the observations, is used to weight the diversity estimate based on probability. The difference resides in that the disparity in this continuous measure is absolute, while in Rao’Q it is relative between two observations.

This difference makes more complex the generalisation to a multi-layer, where this time the uni-dimensional cumulative residual probability is substituted with a multivariate one. For instance, here is an example making use of three layers / bands:

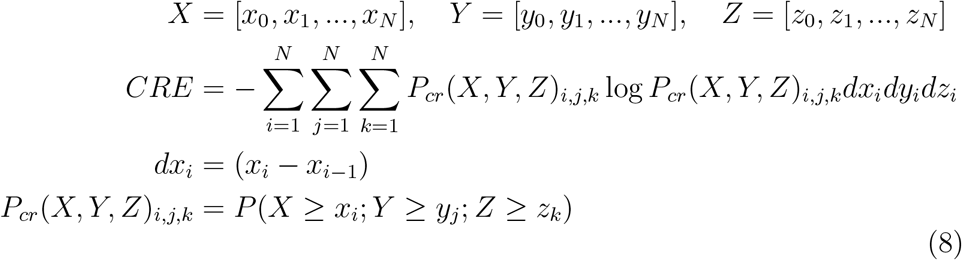

The calculation of the cumulative residual probability *P*_*cr*_(*X, Y, Z*) in an efficient way is based on: i) calculating a contingency array with a certain dimension for each band, and then ii) performing a reverse cumulative sum along each dimension as follows:

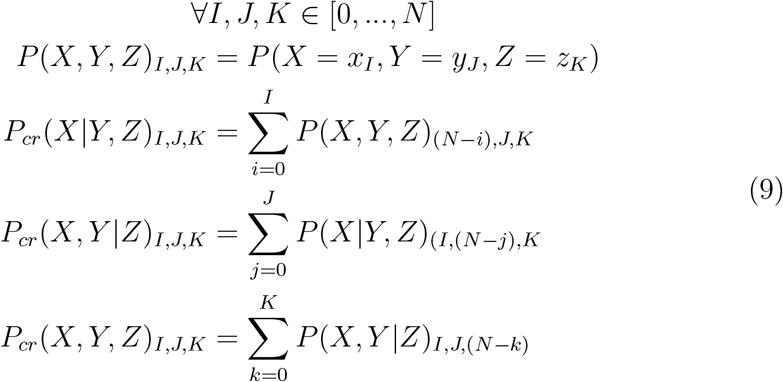

In `rasterdiv` Cumulative Residual Entropy can be calculated as:

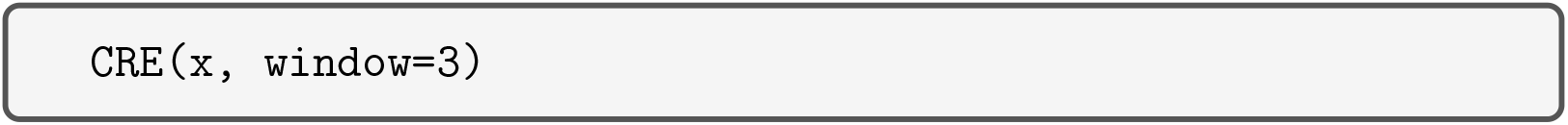

producing a map such as that achieved in Figure 5.

**Figure 5:**
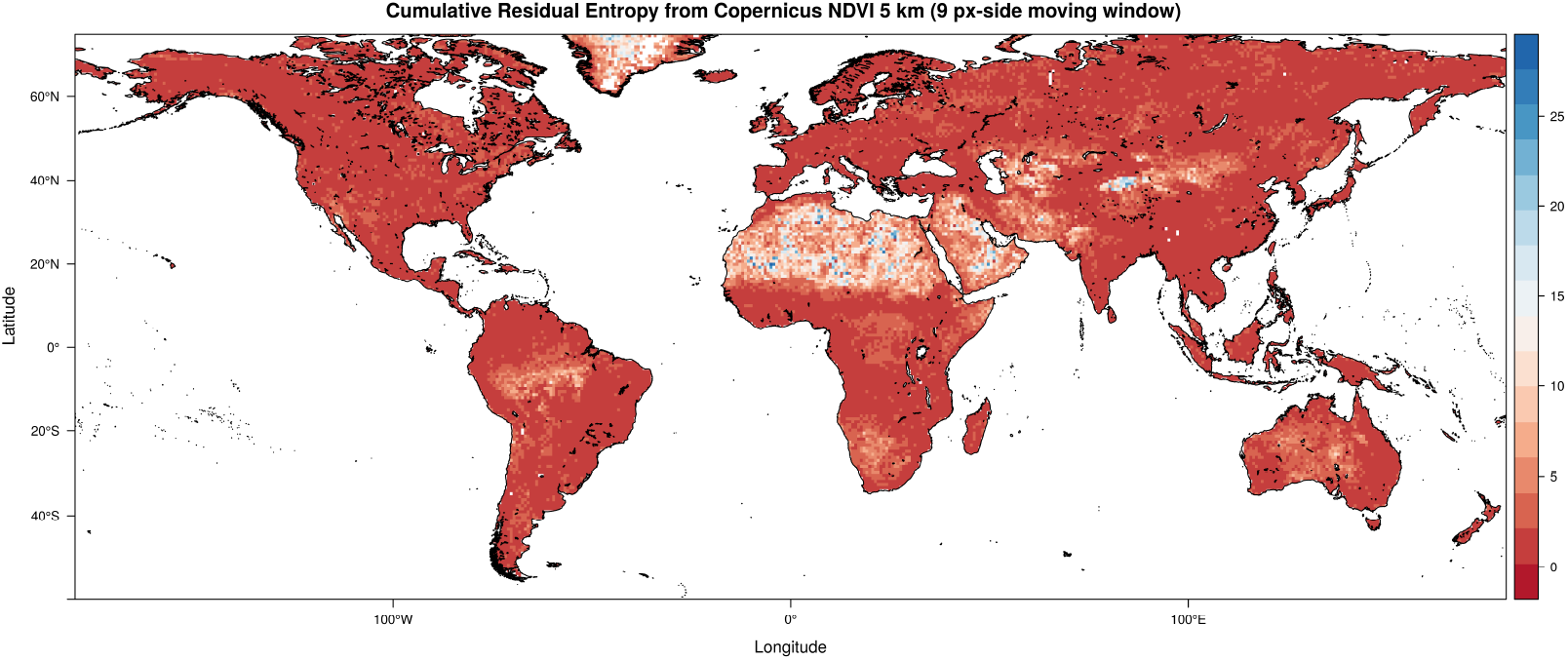
Cumulative Residual Entropy calculated on a Copernicus Proba-V NDVI image at 5km. This figure has been generated by the command `cre <-CRE(ndvi17 r,window=9,np=8,cluster.type=“SOCK”,na.tolerance=0.5)`.

## 5 Solving point descriptors of diversity: the Rényi and Hill generalised entropy

The metrics described above represent point descriptors of diversity, each of which is able to represent only a part of the whole diversity spectrum that can be attained. There is actually no single measure that could be adopted to represent all the different aspects of diversity with an intrinsic fallacy in considering a ‘true’ diversity (Gorelick, 2011a).

Rényi (1970) firstly proposed a measure which is able to represent several diversity metrics in just one formula, by only changing one parameter (*α* in the original version of his manuscript). Given a sample area with *N* pixel values and *p*_*i*_ relative abundances for every *i* ∈ {1, …, *N*}, the Rényi index is:

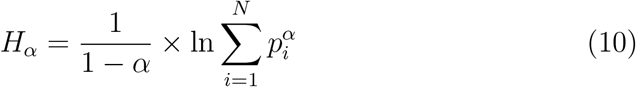

Changing the parameter *α* will lead to different indices starting from the same formula (Hill, 1973). As an example, when *α*=0, *H*_0_ = ln(*N*) where *N* =richness, namely the maximum possible Shannon index (*H*_*max*_). In practice, with *α* = 0, all the spectral values equally contribute to the index, without making use of their relative abundance. For *α* ⟶ 1, the Rényi will equal Shannon *H*, according to the l’Hospital’s rule, while for *α*=2 the Rényi index will equal the ln(1/*D*) where *D* is the Simpson’s dominance (Simpson, 1949). The theoretical curve relating the Rényi index and *α* is a negative exponential, i.e. it decays until flattening for higher values of *α*, where the weight of the most abundant spectral values is higher with small differences among the attained diversity maps (Ricotta et al. (2003a)).

In `rasterdiv` the Rényi index is calculated as:

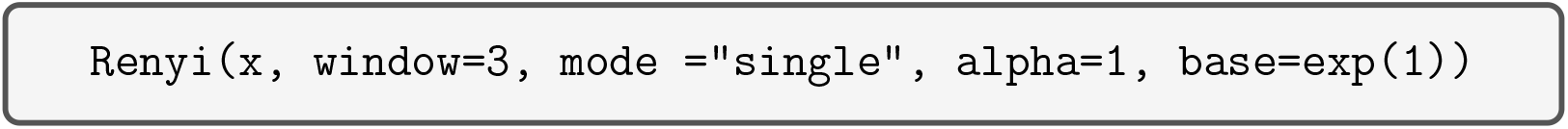

where `x` and `window` are the input set and the window size of analysis, as in previous functions (Figure 6). The mode can be i) `“single”` to compute the Rényi index for just one alpha value, ii) `“iterative”` to compute the Rényi index for all the integer values of alpha in a given interval, or `“sequential”` to compute the Rényi index for all the alpha values in a given vector. `alpha` indicates the *α* vale in Equation 10. Its default value is 1. In `“single”` mode, alpha has to be a numerical value greater than 0; in `“iterative”` mode, alpha has to be a length 2 vector and in `“sequential”` mode, alpha has to be a vector of length at least 2. `base` is a numerical value, which let the user choose the base of the logarithm in Rényi index formula. Its default value is exp(1).

**Figure 6:**
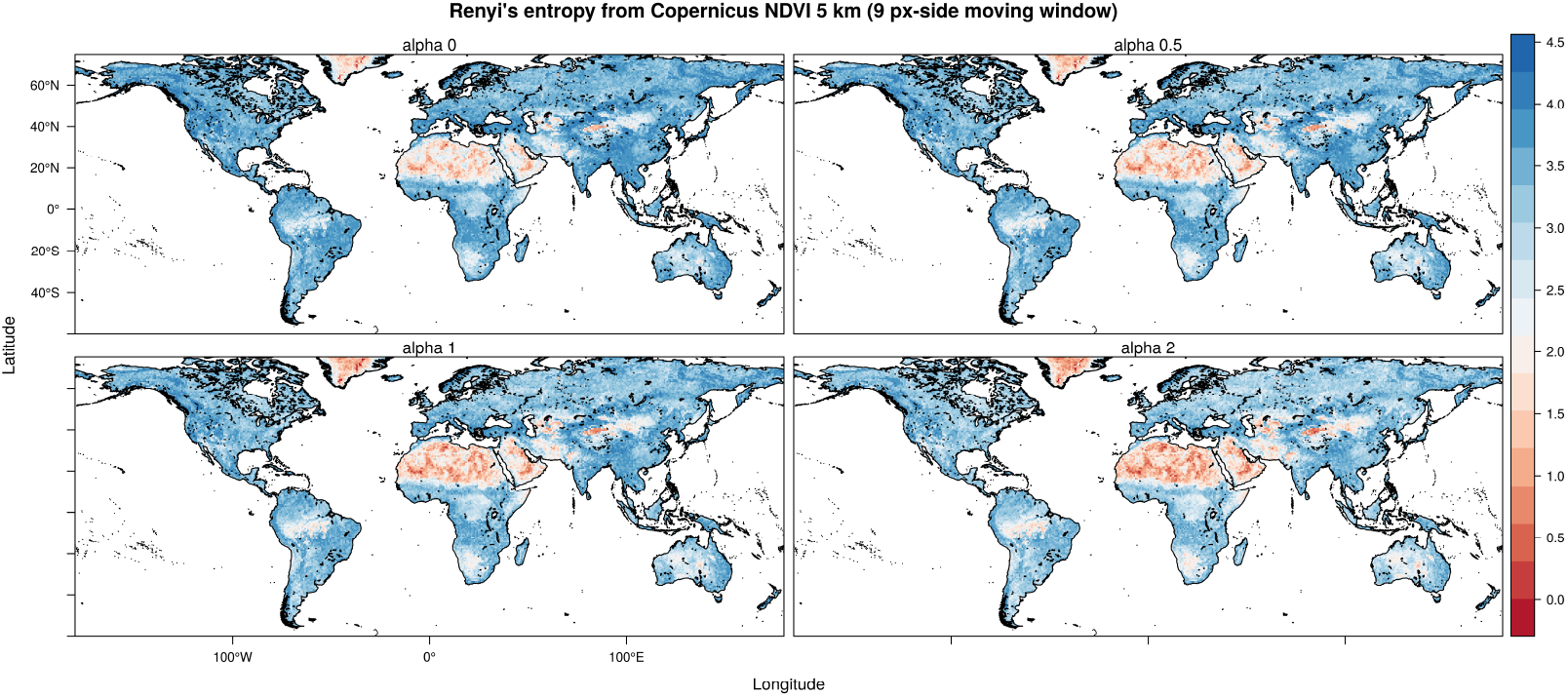
Rényi index calculated on a Copernicus Proba-V NDVI image at 5km, considering different *α* values, from 0 to 2. With *α* → 1, the diversity map is equal to the Shannon’s map of Figure 2. Increasing *α* will create a flattening of the index with a lower ability to discern differences among different maps (see Appendix 2). This figure has been generated by the command `ren <-Renyi(ndvi17 r,window=9,np=8,cluster.type=“SOCK”,alpha=c(0,0.5,1,2))`.

Hill (1973) was the first ecologist applying the generalised entropy concept initially developed by Rényi (1970). In particular, since no particular formula would have a preeminent advantage over the others (Hill, 1973), the Hill’s generalised entropy *N*_*α*_ was based on the effective number of species of *H*_*α*_, namely the number of species that would lead to *H*_*α*_ if they were equally abundant. In our case, the “species concept” is translated to the “spectral values” concept. Hence, *N*_*α*_ is the effective number of spectral values that would give *H*_*α*_ as an output. *N*_*α*_ can thus be calculated as:

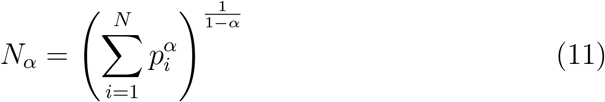

As for the Rényi generalised entropy, changing *α* will let the index transform in many other widely used indices, which are point descriptions of diversity, i.e. peculiar cases of the Hill’s generalised theory. Hence, for *α* = 0, *N*_0_ = *N*, where *N* is the total number of spectral values in the window of analysis; for *α* = 1, *N*_1_ = exp *H*; for *α* = 2, *N*_2_ = 1/*S*, where *S* is the Simpson’s index, and for *α* = ∞, 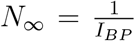, where *I*_*BP*_ is the Berger-Parker index (Figure 7). We refer to Ricotta et al. (2003a) and Ricotta et al. (2003b) for a concise review on the theoretical properties of the Rényi and the Hill’s generalised entropy, respectively. In `rasterdiv`, the Hill’s generalised entropy can be calculated as:

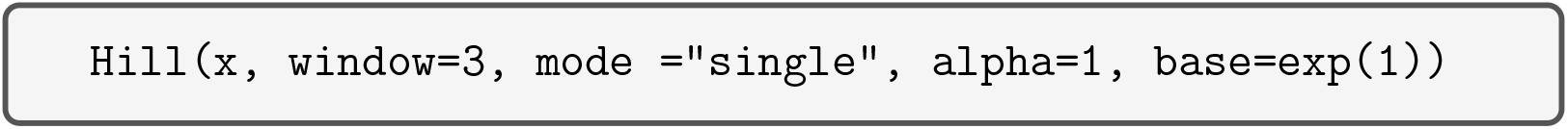

**Figure 7:**
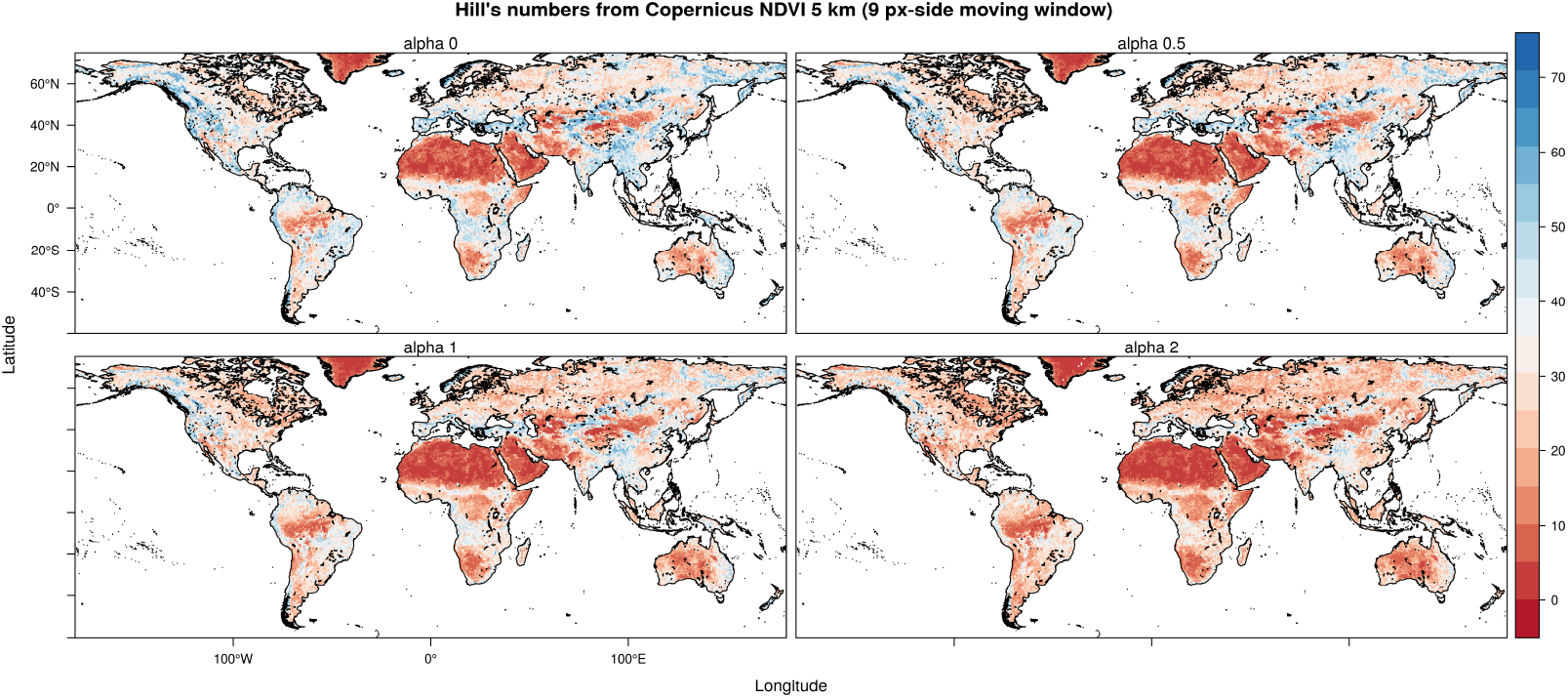
Another generalised entropy measure of diversity: the Hill index, for which the same reasoning of the Rényi index holds true. The maps are derived from a Copernicus Proba-V NDVI image at 5km. This figure has been generated by the command `hil <-Hill(ndvi17 r,window=9,np=8,cluster.type=“SOCK”,alpha=c(0,0.5,1,2))`.

The Hill’s generalised entropy can be computed for i) one *α* value in `single mode`, or ii) all the integer values of *α* in a given interval (`iterative mode`). We refer to Chao et al. (2016) for a complete overview of the Hill’s numbers application in ecology.

## 6 Discussion

In this paper, we provided a full description of the main functionalities of the new `R` package `rasterdiv`. The `rasterdiv` package provides an unprecedented suite of functions to calculate different indicies for estimating diversity from space and to perform a first exploration of potential biodiversity hotspots worldwide at a glance. Of course, measures of diversity from space should not be viewed as a replacement of in-situ data on biological diversity, but they are rather complementary to existing data and approaches. In practice, they integrate available information of Earth surface properties, including aspects of functional (structural, biophysical and biochemical), taxonomic, phylogenetic and genetic diversity (Laliberté et al., 2019).

Obviously, in most of the Information Theory based metrics, only one layer can be used, considering those indices related to relative abundance, apart from the Rao’s Q and the Cumulative Residual Entropy (CRE). In the Rao’s Q index, multidimensional systems can be used to calculate spectral distance (see also Nakamura et al. (2020) on the dimensionality of diversity), while in the CRE it is possible to calculate a multidimensional cumulative distribution to be used in the estimates (Drissi et al., 2008). In general, remotely sensed data are actually the approximation of more complex systems, which depends on the original radiometric and spectral resolution. In ecological terms, such original spectral space formed by many bands is analogous to the Hutchinson’s hypervolume, in which a geometrical order is given to those variables shaping species’ niches (Hutchinson, 1959; Blonder, 2018). In this case, the spectral space is expected to be related to both species niches and their relative diversity. The use of such spaces is an efficient approach to figure out the diversity of an area and potentially guide field sampling and monitoring schemes (Rocchini et al., 2018).

Concerning the data being used, spectral diversity measures computed from satellite images represent a valid alternative to class-based land cover maps for investigating landscapes heterogeneity. For instance, a highly fragmented landscape characterised by a mosaic of crops and seminatural forests suffers from oversimplification when investigated through land cover classes (Amici et al., 2018), while it should present higher spectral diversity values compared to more homogeneous landscapes within the same study area (Rocchini and Ricotta, 2007). Several studies have already acknowledged the importance of computing continuous spectral diversity measures from spectral bands in order to better understand and discriminate the various landscape components (Karlson et al., 2015; Godinho et al., 2018; Ribeiro et al., 2019; Doxa et al., 2020). This said, caution is warranted when making use of continuous data, by seriously considering the radiometry of pixel values. As an example, relying on continuous NDVI (Normalised Difference Vegetation Index) values, ranging from −1 to 1 with float (decimal) precision data, will lead to a high neighbouring diversity which could actually be the effect of data binning rather than of a biological underlying pattern. In general, an 8-bit image with a range of integer values/classes from 0 to 255 would be preferable. In this paper, we made use of an 8-bit NDVI layer rescaled from Copernicus data. However, a multispectral system reduced to one single layer through the first component of a Principal Component Analysis, or similar multidimensionality reduction techniques, would also be useful (Féret and Boissieu, 2020). In fact, NDVI assumes a biomass-grounded reflectance model, while the direct use of the original spectral data (digital numbers) does not generally require any assumptions about the biology of objects being sensed.

As remotely sensed estimates of diversity are currently based on relatively long time series, they might allow a more general forecasting framework of future shifts in rates of diversity change. This is particularly important when aiming at finding potential indicators of diversity change in time (Schmeller et al., 2018). On this point, it has been widely demonstrated that remotely sensed diversity might be in line with most of the spatially-constrained Essential Biodiversity Variables proposed by Skidmore et al. (2015).

The `rasterdiv` package might also be particularly useful when aiming at calculating diversity directly from climate data, derived from remote sensing (Metz et al., 2014). This could allow analysing diversity based on the main drivers of biological diversity in the field, rather than on the patterns resulting from pure spectral response. This is true when considering both wide climatic variations at global scale and microclimate variations at the scale of individuals (Zellweger et al., 2019). Due to unprecedented rates of climatic changes, the adaptation of species to climate change is a benchmark in ecology. Hence, estimating diversity from climate gridded data could improve our understanding of the variability of species ranges at different spatial and temporal scales (Senner et al., 2018).

## 7 Conclusion

Measuring diversity from above and delivering rapid and robust knowledge about diversity over wide regions could be of crucial importance for guiding management practices. From this point of view, the spatial variation of the spectral signal has an intrinsic cumbersome relation with the spatial autocorrelation (sensu Laliberté (2008)) of pixel values over space (and time, e.g. Rocchini et al. (2019)), which renders the proposed `rasterdiv` package a powerful tool to monitor the variation of ecosystems properties over space and time, and thus their change (Rocchini et al., 2018).

As previously stated, no single measure provides a full description of all the different aspects of diversity. That is why, the `rasterdiv` package can result useful in making multiple calculations based on reproducible open source algorithms, robustly rooted on Information Theory from which the different indices are extracted.

## Notes

https://cran.r-project.org/web/packages/rasterdiv/index.html

### Competing Interest Statement

The authors have declared no competing interest.

